# Longevity, clonal relationship and transcriptional program of celiac disease-specific plasma cells

**DOI:** 10.1101/2020.04.30.058560

**Authors:** Ida Lindeman, Chunyan Zhou, Linn M. Eggesbø, Zhichao Miao, Justyna Polak, Knut E. A. Lundin, Jørgen Jahnsen, Shuo-Wang Qiao, Rasmus Iversen, Ludvig M. Sollid

## Abstract

Disease-specific plasma cells (PCs) reactive with transglutaminase 2 (TG2) or deamidated gluten peptides (DGP) are abundant in celiac disease (CeD) gut lesions. Their contribution toward CeD pathogenesis is unclear. We assessed expression of markers associated with PC longevity in 15 untreated and 26 treated CeD patients in addition to 13 non-CeD controls, and performed RNA-sequencing with clonal inference and transcriptomic analysis of 3251 single PCs. We observed antigen-dependent V-gene selection and stereotypic antibodies. Generation of recombinant DGP-specific antibodies revealed a key role of a heavy-chain residue that displays polymorphism, suggesting that immunoglobulin gene polymorphisms may influence CeD-specific antibody responses. We identified transcriptional differences between CeD-specific vs non-disease-specific PCs and between short-lived vs long-lived PCs. The short-lived CD19^+^CD45^+^ phenotype dominated in untreated and short-term-treated CeD, in particular among disease-specific PCs but also in the general PC population. Thus, the disease lesion of untreated CeD is characterized by massive accumulation of short-lived PCs that are not only directed against disease-specific antigens.

## INTRODUCTION

A feature of autoimmune diseases is the secretion of specific autoantibodies by terminally differentiated B cells, plasma cells (PCs). IgA- and IgM-secreting PCs are abundant in the lamina propria in the gut, where antibodies are secreted into and coat the intestinal lumen. Traditionally, gut PCs were thought to be short-lived antibody factories, and other potential roles of PCs in health and autoimmune diseases have not been under much investigation. However, expression of human leukocyte antigen (HLA) class II molecules co-stimulatory molecules such as CD40, CD80 and CD86, as well as cytokine secretion, suggest that PCs can have additional functions.^1-4^ Recently, three distinct subsets of short-, intermediate- and long-lived lamina propria PCs were described based on expression of the cell surface markers CD19 and CD45.^5^ The longevity profile of human gut PCs in a disease setting is yet uncharacterized.

Celiac disease (CeD) is associated with a highly specific autoantibody response. The patients have autoantibodies to the enzyme transglutaminase 2 (TG2) as well as antibodies to deamidated gluten peptides (DGP).^6^ The disease is driven by an immune response to cereal gluten proteins which explains the presence of the DGP antibodies. TG2 is involved in the pathogenesis by catalyzing deamidation of gluten peptides. The patients have CD4^+^ T cells that recognize DGP bound to disease-associated HLA-DQ molecules.^7^ These T cells likely provide T-cell help to DGP-specific B cells as well as TG2-specific B cells; the latter by involvement of TG2-gluten peptide hapten-carrier like molecules.^6^ The disease lesion in the small intestine is characterized by blunting of villi and infiltration of inflammatory cells, including PCs. The active lesion typically has a 2-3 fold increased density of IgA and IgM PCs in lamina propria,^8^ with about 10% of the PCs being specific for TG2 and 1% being specific for DGP.^9,10^ The only available treatment for CeD is adherence to a strict gluten-free diet (GFD). After commencing a GFD, the concentration of disease-specific serum antibodies,^11^ and number of disease-specific PCs in lamina propria drop rapidly within months.^12^

The role of PCs in CeD is currently unknown, and it is uncertain to what extent the secreted autoantibodies play a pathogenic role.^13,14^ Recent reports have shown that intestinal PCs may have additional functions to antibody secretion such as antigen presentation and cytokine secretion,^4,15^ leading to the hypothesis that disease-specific PCs are playing an active role in the pathogenesis of CeD including having a role as antigen-presenting cells (APCs) for gluten-specific T cells. Identifying functions of gut PCs could provide more knowledge about disease mechanisms and point out disease-specific PCs as therapeutic targets.

High-throughput sequencing technologies have revolutionized the profiling of immune receptor repertoires in health and disease (AIRR-seq),^16,17^ and we have previously applied these technologies to dissect the repertoire of disease-specific PCs in CeD.^10,18-22^ However, little is known about the transcriptional state of these cells. The development of single-cell RNA-seq (scRNA-seq) protocols provides a unique tool to identify and characterize cell subsets according to their full transcriptional profile.^23-25^ Additionally, recent computational tools for reconstruction of antigen receptor sequences from scRNA-seq data^26-33^ now allow us to link the transcriptional profile of individual PCs with information about their B-cell receptor (BCR) and thus clonal relatedness.

Here we used scRNA-seq and BCR reconstruction and clonotype inference by the computational tool BraCeR^30^ to characterize PCs in the healthy small intestine and in an autoimmune setting, specifically CeD. We observed antigen-dependent V-gene selection, and we identified a novel stereotypic DGP-specific antibody. We further observed that disease-specific gut PCs in celiac lesions of untreated (UCeD) or short-term treated (TCeD) patients rarely carried a phenotype previously shown to be associated with long life span,^5^ whereas PCs of this phenotype were more abundant in TCeD patients treated for long term. PCs of different longevity phenotype differed in their transcriptomic profiles. Differentially expressed genes included genes involved in metabolism and regulation of apoptosis. The dominance of the short-lived CD19^+^CD45^+^ phenotype in UCeD and short-term-treated CeD, both among disease-specific PCs but also in the general PC population, suggests that new and short-lived PCs massively accumulate in the UCeD lesion.

## MATERIALS AND METHODS

### Ethics statement

This study was approved by the Regional Ethics Committee of South-Eastern Norway (REK 6544). Informed, written consent was obtained from all patients prior to sample collection.

### Sample collection

We collected intestinal biopsies from 15 patients with untreated CeD (UCeD), 26 CeD patients on GFD for minimum one year (TCeD) and 13 non-CeD controls (no known autoimmune diseases or intestinal inflammation) coming in for gastroduodenoscopy at the Department of Gastroenterology, Oslo University Hospital Rikshospitalet (OUH), Norway or Department of Gastroenterology, Akershus University Hospital (AUH), Norway. CeD diagnosis was given following the British Society of Gastroenterology guidelines.^34^ Pathological and laboratory data (Suppl. Table 1) are taken from the hospital records. Duodenal biopsies were collected in ice-cold RPMI 1640 medium (samples from OUH) or phosphate-buffered saline (PBS) (samples from AUH). Samples from all individuals were used for flow cytometry analysis, while four UCeD patients, three TCeD patients and five controls also were analyzed by scRNA-seq.

### Sample processing

Intestinal biopsies were treated twice with 2 mM EDTA in PBS with 2% fetal calf serum (FCS) at 37°C under constant rotation for 10 min in order to remove the epithelium and intraepithelial lymphocytes. The biopsies were subsequently digested with 1 mg/ml collagenase type H (Sigma) in 2% FCS in PBS at 37°C for 35 min under constant rotation. The digested tissue was homogenized with a syringe and needle and filtered through a 40-μm cell strainer and washed with PBS. Samples were then immediately used for analysis and/or sorting. Sixteen of the samples had been cryopreserved before flow cytometry analysis (Suppl. Table 1).

### Fluorescence-activated cell sorting (FACS)

We used fluorescently labeled tetramers loaded with TG2 and deamidated gluten peptides (DGP) in order to identify CeD-specific cells. BirA-tagged TG2 was produced in Sf9 insect cells, and N-terminally biotinylated.^35^ The immunodominant deamidated gluten epitope PLQPEQPFP (DGP) was obtained as biotinylated peptide (GL Biochem, Shanghai). TG2 and DGP were multimerized on PE-labeled streptavidin (SA) (Invitrogen) and APC-SA (PhycoLink), respectively, on ice for at least 45 min at a 4:1 ratio of TG2/DGP:PE/APC-SA. The single cell suspension was stained with TG2-SA-PE, DGP-SA-APC, and the following anti-human antibodies: CD3-BV605 (UCHT1, Biolegend) or CD3-Super Bright 600 (UCHT1, eBioscience), CD11c-BV605 (3.9, Biolegend), CD14-BV605 (M5E2, Biolegend), CD45-APC-Cy7 (H130, Biolegend), CD19-Pacific Blue (HIB19, Biolegend) and CD38-FITC (HB7, eBioscience) in PBS with 2% FCS for 30 min on ice, washed with 2% FCS PBS and filtered before cell sorting using a BD FACSAria II (OUH Radiumhospitalet) or BD FACSAria III instrument (OUH Rikshospitalet) with FACS Diva software. Flow cytometry data were analyzed with FlowJo (BD) v10.

Single, large CD38^++^CD3^-^CD11c^-^CD14^-^ PCs were randomly index-sorted according to antigen specificity (Suppl. Fig. 1C). We aimed at obtaining approximately equal numbers of TG2-specific PCs and PCs of unknown specificity on each plate. DGP-specific PCs, being rare, were also selectively sorted for some patients. For two of the controls, we sorted PCs based on their CD19/CD45 phenotype as a proxy for cell longevity (Suppl. Fig. 1D). Cells were sorted into 96-well plates containing 2 μl lysis buffer per well (0.2% v/v Triton X-100 (Sigma) in H_2_O with 2 U/μl RNase inhibitor), immediately spun down after sorting and placed on dry ice before being stored at −70°C until further processing. Cells of different specificities and longevities were sorted on the same plate to minimize batch effects.

**FIGURE 1.**
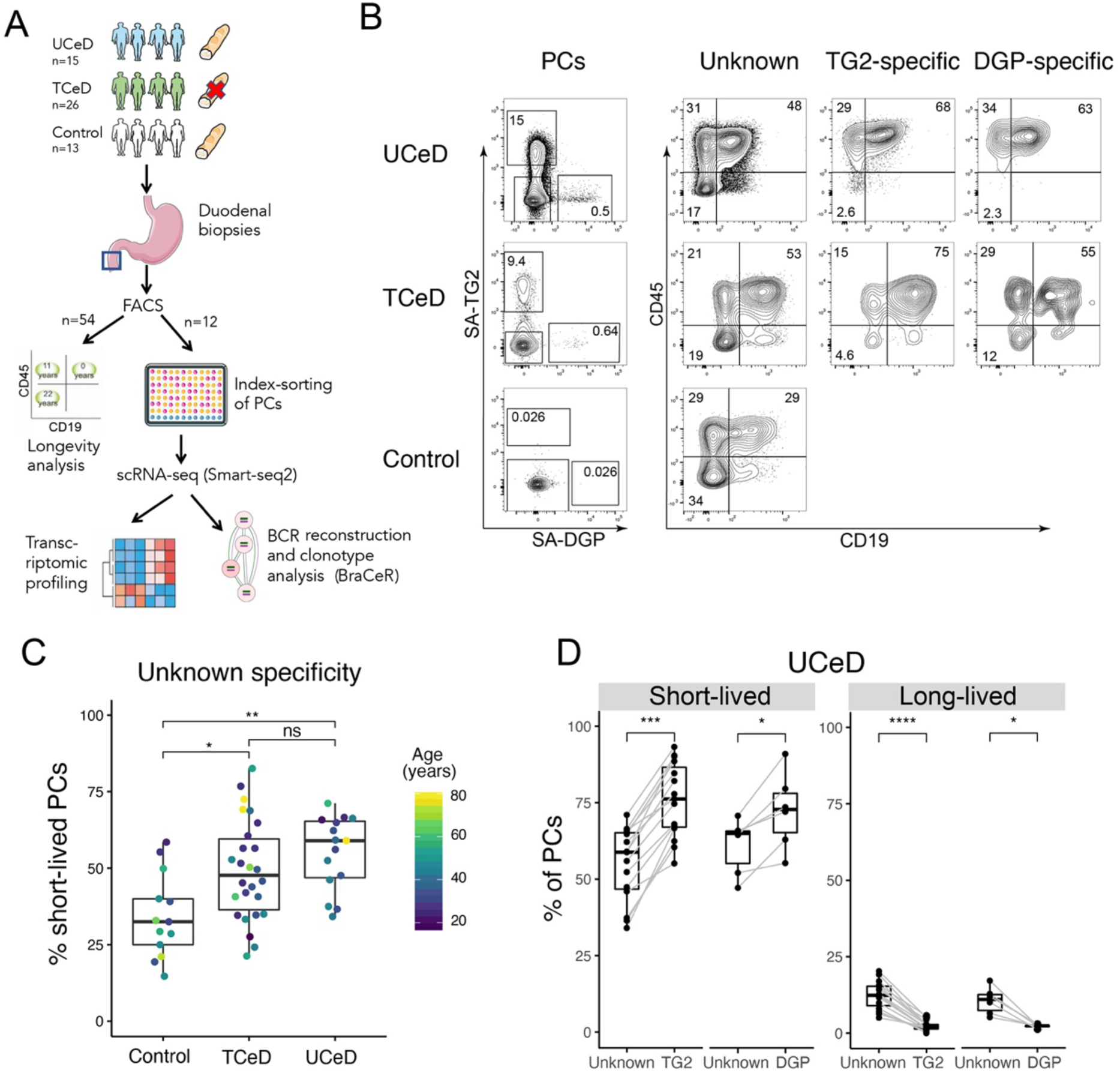
Analysis of PC longevity by flow cytometry. **A.** Experimental approach. UCeD = untreated celiac disease (CeD), TCeD = treated CeD, PC = plasma cell, BCR = B-cell receptor, FACS = fluorescence-activated cell sorting. The red cross indicates a gluten-free diet. The blue square indicates location of biopsy sampling (duodenum). **B.** Representative flow cytometry data showing percent TG2-specific and DGP-specific PCs and the longevity markers CD19/CD45 for each specificity for one UCeD, one TCeD and one control subject. PCs were gated as large, single CD3-CD11c-CD14-CD38++ cells. **C.** Percent of PCs of unknown specificity belonging to the short-lived CD19+CD45+ population in UCeD (n=15), TCeD (n=26) and controls (n=13). *p*-values were calculated with an unpaired Wilcoxon rank-sum test (*=*p*<.05, **= *p*<.01). **D.** Summarized data for longevity in UCeD (n=15) patients. Only patients with at least 30 recorded DGP-specific PCs are included in the comparisons between DGP-specific PCs and PCs of unknown specificity (n=6). *p*-values were calculated with a paired Wilcoxon signed-rank test (*=*p*<.05, ***= *p*<.001, ****=*p*<.0001).

### Single-cell RNA-sequencing (scRNA-seq)

We performed scRNA-seq of in total 3743 intestinal PCs. The cells were subjected to reverse transcription and cDNA pre-amplification according to a modified version of the Smart-seq2 protocol,^36^ using 0.5 μl/well SMARTScribe Reverse Transcriptase (Clontech) for reverse transcription. The cDNA was pre-amplified with 21 PCR cycles and purified with 20 μl Ampure XP beads per well (Agencourt). Amplified cDNA was tagmented using an inhouse-produced Tn5 transposase^37^ and dual indexed with Nextera (XT) N7xx and S5xx index primers at a final concentration of 125 nM. Four plates were pooled together and sequenced at the Norwegian Sequencing Centre on a NextSeq500 instrument with 75 bp paired-end reads in high-output mode. Average number of read pairs per cell was around one million.

### scRNA-seq quality control

Raw reads were trimmed for adapter sequences and low-quality sequences using Cutadapt v.1.18^38^ through Trim Galore v0.6.1 in paired-end mode. We quantified transcript expression for the scRNA-seq data with Salmon v0.11.3^39^ using cDNA sequences from GRCh38.94 and a k-mer length of 25 for creating the salmon index. Transcripts were aggregated to gene level and a transcript-length-corrected gene expression matrix for all the cells was constructed using tximport v1.8.0.^40^ The single cells were subsequently filtered based on the following quality measures to discard low-quality cells: Number of detected genes, number of reads, percent mitochondrial genes, percent reads mapping to the reference transcriptome, percent immunoglobulin (Ig) genes and whether a BCR heavy chain could be reconstructed by BraCeR. Library-specific quality threshold values are shown in Supplementary Table 3. Quality control was performed in R version 3.5.3 using the scater package.^41^ In total 3251 cells remained after quality control.

### scRNA-seq batch correction and normalization

Since a large part of the PC transcriptome (67.7±8.4% in our data) consists of Ig genes and these genes would dominate downstream analyses and mask non-Ig-related transcriptional differences between cell populations, all Ig genes were discarded from the analysis before normalization. After removing Ig genes, genes with more than three reads detected in more than 20 cells were retained. For each analysis, the gene expression matrix was subset according to the cells of interest and normalized by total counts per cell using scanpy v1.4.4^42^ in python 3.7. The normalized expression matrix was then logarithmized as *X* = ln(*X* + 1). Highly variable genes were identified with the *highly_variable_genes* function of scanpy with the following parameters: *min_mean=0.1, max_mean=10, min_disp=0.25*. Unwanted variation, such as the number of genes and reads detected and percent mitochondrial genes, was then regressed out using NaiveDE v1.2.0 (https://github.com/Teichlab/NaiveDE) and the highly variable genes, while retaining the variation of interest for each analysis (see figure legends for analysis-specific details). The expression matrix was scaled using sklearn (scikit-learn v0.21.3).^43^

### PCA and UMAP

We performed Principal Component Analysis (PCA) of the scaled expression matrices using scanpy, and investigated which variables drove the different principal components. The principal components explaining most of the variation in the variable of interest were then used as input for Uniform Manifold Approximation and Projection for Dimension Reduction (UMAP).^44^

### Identifying differentially expressed genes

Differentially expressed genes were identified with the scanpy function *rank_genes_groups* using the corrected gene expression matrix as input and Wilcoxon ranked-sum test, correcting for multiple testing with the Benjamini-Hochberg method. The differentially expressed genes were then filtered with *filter_rank_genes_groups* with the following parameters: *min_fold_change* (specific values are stated in figure legends), *min_in_group_fraction=0.3, max_out_group_fraction=1*. Only genes with an adjusted *p*-value < 0.05 were retained. All groups were tested against the remaining cells unless otherwise stated.

### BCR reconstruction with BraCeR

Paired BCR heavy and light chains were reconstructed for each cell using BraCeR in assembly mode with all raw reads as input. BraCeR assembly was run with *--threshold 5000* in order to filter out lowly expressed reconstructed BCRs that may arise from contamination between wells or errors in indices. Clonally related PCs were determined separately for each patient and each PC specificity for cells passing scRNA-seq quality control using BraCeR in summary mode with the following parameters: *--include_multiplets --infer_lineage*. Potential cell multiplets were manually inspected and removed as previously outlined.^45^ Somatic mutations in the variable regions were analyzed with IMGT/HighV-QUEST,^46^ and SHM rates were calculated by dividing the number of point mutations by sequence length.

### Expression of monoclonal antibodies, site-directed mutagenesis and ELISA

BCR sequences obtained from two DGP-reactive PCs using the *IGHV3-74*:*IGKV4-1* gene pair were expressed as recombinant human IgG1 molecules as previously described.^47^ Briefly, synthetic DNA (GenScript) encoding the antibody variable regions with either R or A in IMGT position 55 of the heavy chain were cloned into expression vectors and used for transfection of HEK 293-F cells. The secreted antibodies were subsequently purified from the supernatant on HiTrap Protein G columns (GE Healthcare). Plasmid DNA encoding a gliadin-reactive *IGHV4-4*:*IGKV4-1* antibody with R in position 55 (corresponding to the *IGHV4-4*07* allele) was already available in the lab.^10^ In this case, an R to E (found in alleles *\*01* to *\*06*) substitution was introduced by overlap extension PCR. The reactivity of all antibody variants with biotinylated, synthetic DGP was assessed by ELISA. Briefly, the peptide (50 nM) was immobilized in SA-coated microplates (Thermo Scientific) followed by incubation with different concentrations of recombinant antibodies in TBS supplemented with 0.1% (v/v) Tween20. Bound antibody was detected with alkaline phosphatase-conjugated goat anti-human IgG (Southern Biotech).

### Statistical analysis

Differences between population distributions were tested using a paired Wilcoxon signed-rank test for data in which TG2-specific, DGP-specific and PCs of unknown specificity were compared against each other within each patient. In order to test differences in distribution between unpaired samples, such as between CeD patients and controls, an unpaired Wilcoxon rank-sum test was used. Results of the statistical tests are throughout this paper given as *p-*value significance levels (*=*p*<.05, **=*p*<.01, ***= *p*<.001, ****=*p*<.0001) after adjusting for multiple testing. Global differences between means were calculated with one-way ANOVA and reported in addition to individual adjusted *p*-values. *p-*values were calculated and added to plots using ggpubr v0.2.4. Linear regression was performed in R with the lm method.

## RESULTS

### Study participants

Information about the study participants is presented in Supplementary Table 1. Subjects in the TCeD group had been on a gluten-free diet (GFD) for 1-31 years (median 2.25 years). Non-CeD controls had no known autoimmune diseases or intestinal inflammation. The median ages of the participants were 38 (UCeD), 34.5 (TCeD), and 47 years (controls) (Suppl. Fig. 1A).

### TG2-specific and DGP-specific gut PCs have a short-or intermediate-lived phenotype in UCeD and short-term TCeD

Using flow cytometry, we found that 17.0±8.7% (mean±sd) of the intestinal PCs from UCeD patients were TG2-specific and 0.48±0.25% were specific for DGP (Suppl. Fig. 1B and Suppl. Table 2). TCeD patients had 6.2±5.7% TG2-specific and 0.23±0.22% DGP-specific PCs persisting after minimum a year on GFD, with decreasing populations of TG2- and DGP-specific PCs in more long-term TCeD patients (Suppl. Fig. 1C). The controls had negligible frequencies of TG2-specific (0.08±0.09%) and DGP-specific (0.02±0.03%) PCs.

In order to assess the longevity of PCs in the gut of CeD patients and non-CeD controls, we analyzed the surface expression of CD45 and CD19 (Fig. 1 and Suppl. Table 2). TG2-specific and DGP-specific PCs from CeD patients largely belonged to the short-lived CD45^+^CD19^+^ or intermediate-lived CD45^+^CD19^-^ compartments (Fig. 1B). Significantly more CeD-specific PCs in UCeD patients belonged to the short-lived compartment compared to PCs of unknown specificity from the same patients (Fig. 1B and 1D). Interestingly, even when looking only at PCs of unknown specificity, UCeD patients had a significantly higher percentage of short-lived PCs compared to the controls (Fig. 1C).

Next, we investigated whether the distribution between short-, intermediate- and long-lived PCs in CeD patients changes with the duration of treatment (years on GFD) (Fig. 2). The percentage of short-lived PCs decreased with time on GFD for both TG2- and DGP-specific PC, while no such trend was observed for PCs of unknown specificity (Fig 2A-B). Conversely, the percentage of long-lived PCs among the TG2- and DGP-specific cells increased with time on GFD. By grouping the TCeD patients according to treatment duration, we found that short-term TCeD patients (treated up to 7.5 years) had a composition of PC longevities similar to UCeD patients, in that the TG2-specific and to some extent the DGP-specific PCs were enriched for short-lived and depleted of long-lived PCs (Fig. 2). We observed the opposite trend for long-term TCeD patients (treated for 9-31 years), which showed a marked decrease in short-lived and increase in long- and intermediate-lived PCs among CeD-specific PCs compared to PCs of unknown specificity (Fig 2C-D).

**FIGURE 2.**
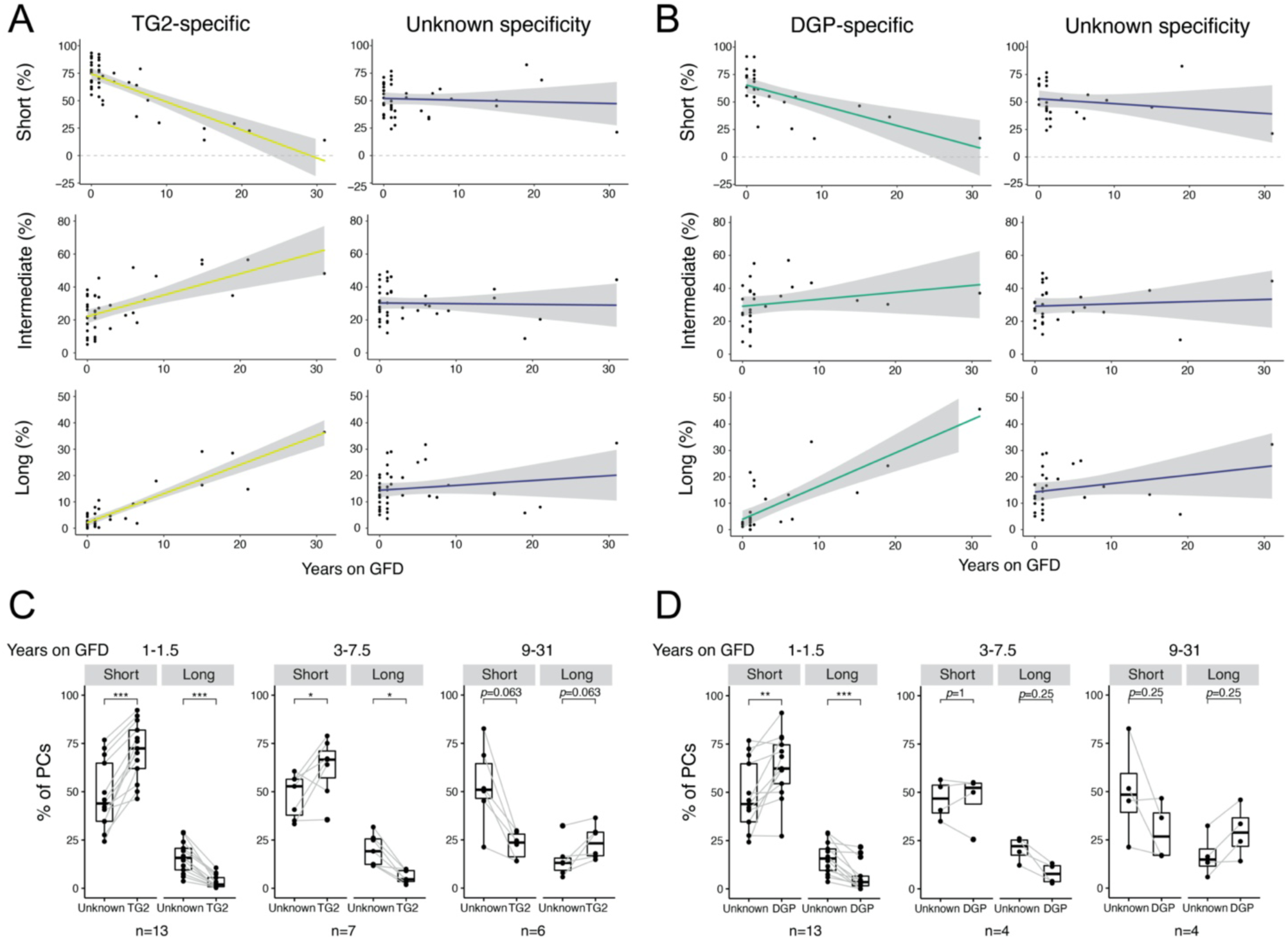
Analysis of PC longevity in TCeD patients using flow cytometry. **A, B.** Percent of PCs belonging to the short-lived (CD19+CD45+), intermediate-lived (CD19+CD45-) and long-lived (CD19-CD45-) compartments for TG2-specific (A) or DGP-specific (B) PCs (left panel) and PCs of unknown specificity (right panel) as a function of treatment duration (years on GFD). UCeD patients are plotted as 0 years on GFD. The blue line represents fitted trend using linear regression. The grey shade indicates the 95% confidence interval of the fitted line. All CeD patients are included in (A) (n = 41), while only patients with at least 30 recorded DGP-specific PCs are included in (B) (n= 27). **C, D.** Percent of PCs belonging to the short-lived and long-lived compartments for TG2-specific (C) or DGP-specific (D) PCs compared to PCs of unknown specificity. Data points from the same patients are connected with grey lines. TCeD patients are grouped according to treatment length. Only patients with at least 30 recorded DGP-specific PCs are included in (D). *p*-values were calculated with a paired Wilcoxon signed-rank test (*=*p*<.05, **=*p*<.01, ***= *p*<.001).

Taken together, our data demonstrate that CeD-specific PCs are skewed toward a short-lived phenotype in UCeD and short-term TCeD patients, and suggest that intermediate- and long-lived PCs are relatively more prevalent within the CeD-specific PC populations with time on GFD.

### Intestinal PCs express mRNA transcripts for certain HLA class II molecules, co-stimulatory molecules and cytokines

A recent bulk RNA-seq study from our group revealed antibody-independent functions of intestinal PCs in health and in CeD.^15^ However, in this study contamination from other cell types could not be completely ruled out as only mean gene expression within populations was assessed. In order to confirm previous findings and further investigate the transcriptional profiles of intestinal PCs at a single-cell resolution, we performed scRNA-seq of intestinal PCs from CeD patients and controls (Fig. 3 and Suppl. Fig. 1). A total of 3251 PCs were analyzed for clonal inference and transcriptional profile. We first looked for expression of genes associated with antigen presentation, and detected expression of *CD74* and HLA class II genes, in particular *HLA-DQA1*, in a large part of the cells regardless of PC specificity, longevity or disease status. Notably, we also detected expression of the co-stimulatory molecules CD40, CD48 and to a small extent CD86, but not CD80, in a population of the PCs. Additionally, *ICAM1, ICAM2* and *ICAM3* transcripts were widely expressed.

**FIGURE 3.**
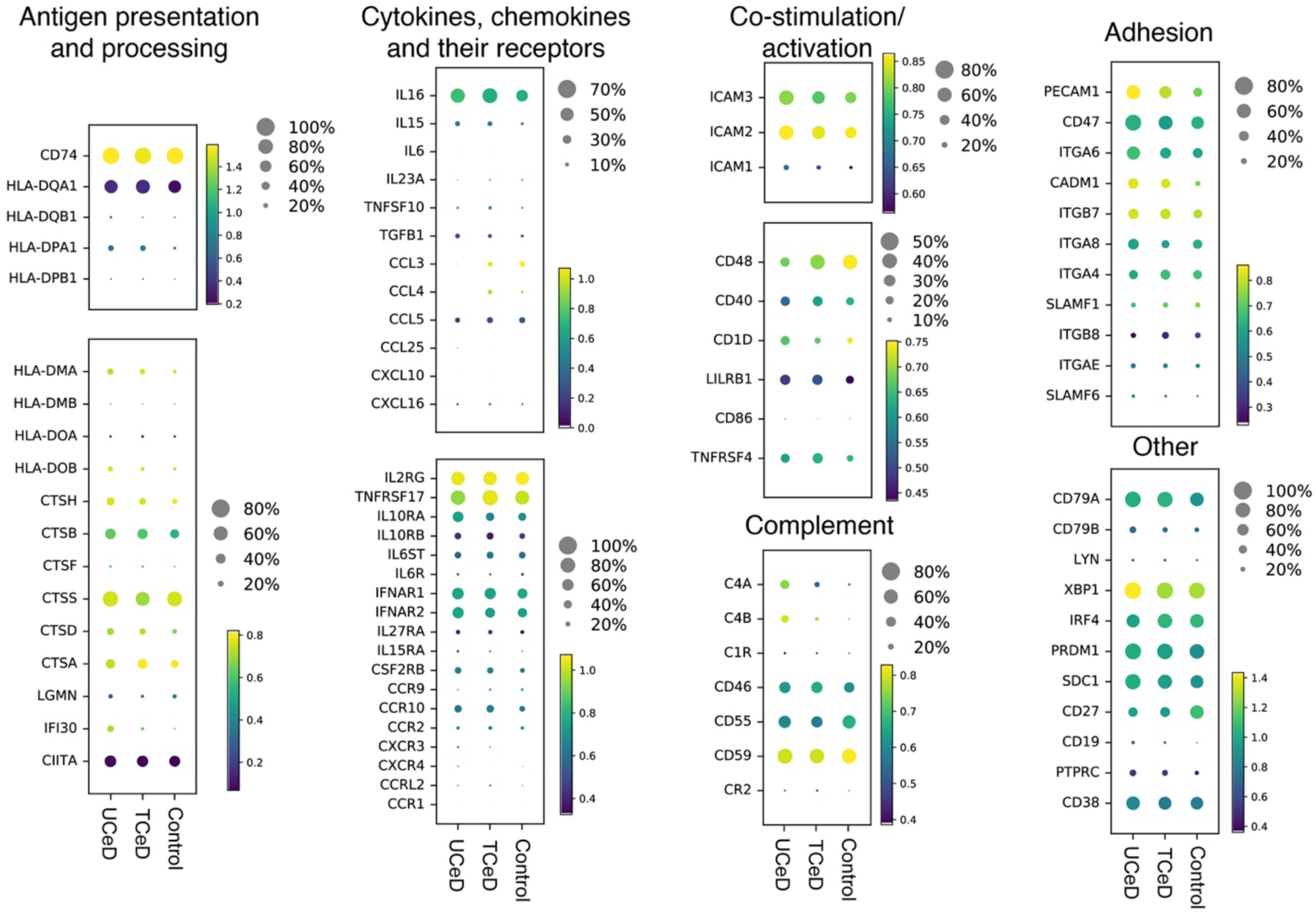
Transcriptional profiling of intestinal PCs using scRNA-seq. Scaled expression data for PCs from all individuals were pooled together according to disease status (UCeD (n=4), TCeD (n=3), control (n=5)). Genes of particular interest are shown in dotplots. Colors represent mean expression of each gene among cells in which the gene was expressed. Dot size represents proportion of cells expressing each gene.

Next, we looked for evidence of cytokine and/or chemokine secretion. Indeed, we detected expression of mRNA transcripts for interleukin (IL)-15, IL-16 and/or TGF-β1 in a substantial population of cells in all patients and controls. Expression of the chemokine genes *CCL3, CCL4* and *CXCL16* was also detected in some of the PCs. Finally, we found wide expression of genes encoding various cytokine/chemokine receptors, such as *TNFRSF17* (BCMA) and *IL2RG*. Our transcriptional analysis of intestinal PCs thus supports the notion that PCs have additional functions to antibody secretion.

### Short- and long-lived intestinal PCs differ transcriptionally

Having looked at the general transcriptional phenotype of intestinal PCs, we next investigated transcriptional differences between short-, intermediate- and long-lived PCs. As few long-lived PCs were index-sorted by FACS for the first three controls, we strictly gated on each longevity population and sorted equal numbers of these from two additional controls in order to compare these clearly defined subsets (Suppl. Fig. 1E).

scRNA-seq followed by principal component analysis (PCA) and UMAP revealed that short-lived PCs to some extent differ transcriptionally from long- and intermediate-lived PCs (Fig. 4A). In order to identify which genes explain the transcriptional differences between the cell populations, we performed differential expression analysis. Our analysis revealed that *CD27* was differentially expressed between all three subpopulations, with highest expression in long-lived PCs and lowest expression in short-lived PCs (Fig. 4B-C). Among the genes that were more highly expressed in long-lived compared to short-lived PCs, we found genes involved in antigen presentation on MHC class I (*HLA-A, HLA-B*, and *HLA-C*), metabolism (*AMPD1* and *QPCT*), inhibition of apoptosis (*BIRC3)*, and *CD9*. Genes that were overexpressed in the short-lived PCs were heavily enriched for genes encoding ribosomal proteins and genes involved in metabolism (*PDIA4, UBE2J1, SPCS1, UBB, TRIB1, ATP5F1A*), possibly reflecting higher antibody secretion in the short-lived PCs.^5^ As expected, *CD19* was exclusively detected in short-lived (CD19^+^) PCs, while *PTPRC* (CD45) was enriched in the short- and intermediate-lived subsets (CD45^+^) (data not shown). The highest ranking differentially expressed genes were also identified when looking at each patient individually. Thus, we demonstrate that PCs of different longevities exhibit distinct transcriptional profiles.

**FIGURE 4.**
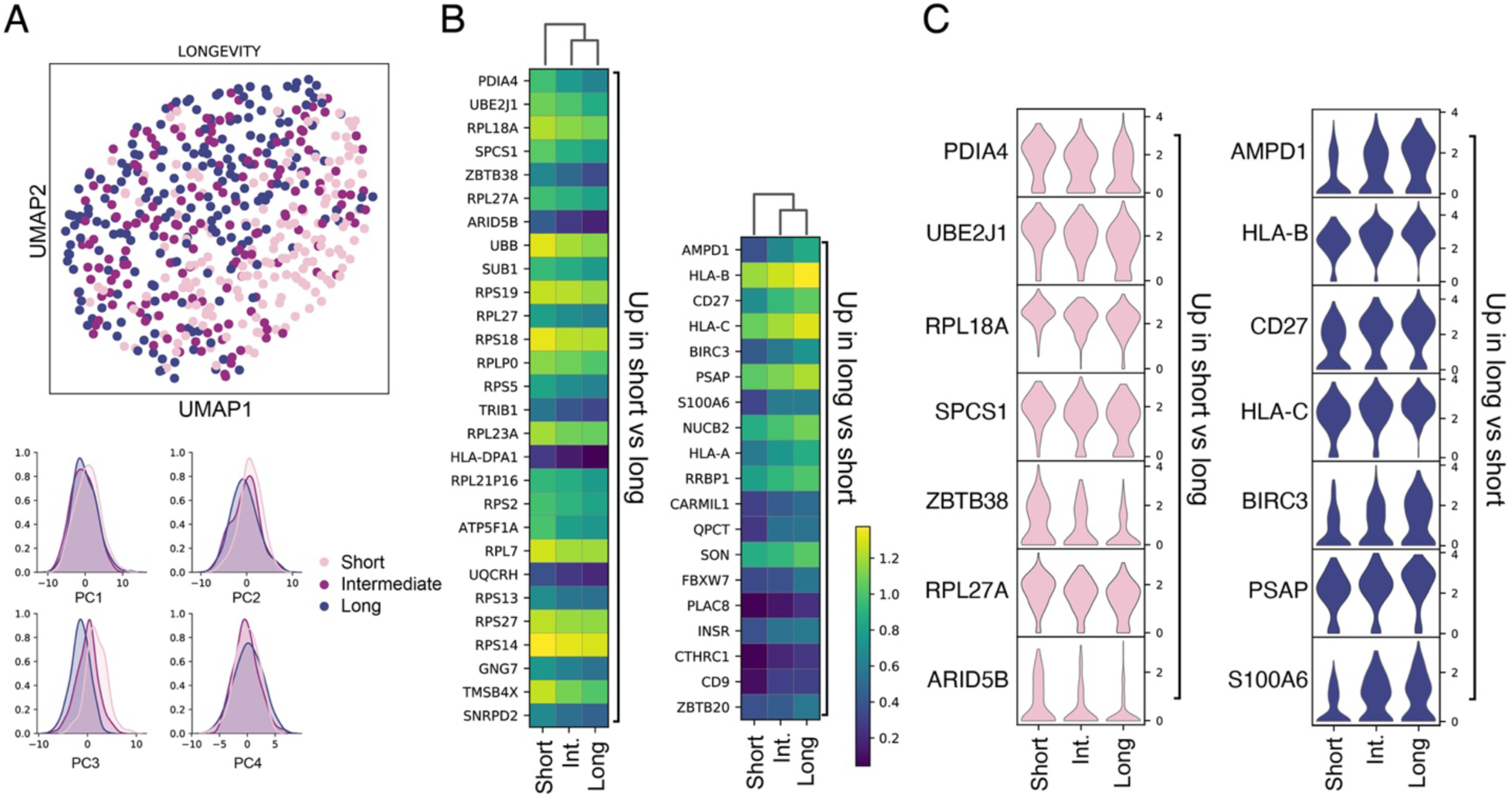
Transcriptional differences between PC longevities. The analysis was performed pooling all PCs from two controls (CD2158 and CD2161) for which equal numbers of cells were sorted for each longevity phenotype. Variation explained by sequencing library or patient was regressed out, while retaining variation explained by PC longevity. **A.** UMAP projection based on the first five principal components (top) and individual principal components, showing variation explained by longevity. **B.** Heat maps showing all statistically significant differentially expressed genes between short-lived and long-lived PCs after filtering. Filtering was done with *min_fold_change=1.2*. **C.** Violin plots showing the top seven upregulated genes in short-lived (left) and long-lived (right) PCs.

### TG2-specific gut PCs have characteristic transcriptional features

Next, we set out to investigate whether TG2-specific or DGP-specific PCs are transcriptionally distinct from PCs of unknown specificity in CeD. When looking at the overall transcriptional landscape of the PCs by dimension reduction with PCA and UMAP, TG2-specific and DGP-specific PCs occupied a slightly different space compared to PCs of unknown specificity with the TG2-specific cells having the most distinct profile (Fig. 5A). As for the controls, we also found PC longevity to be a driver of transcriptional heterogeneity in CeD patients.

**FIGURE 5.**
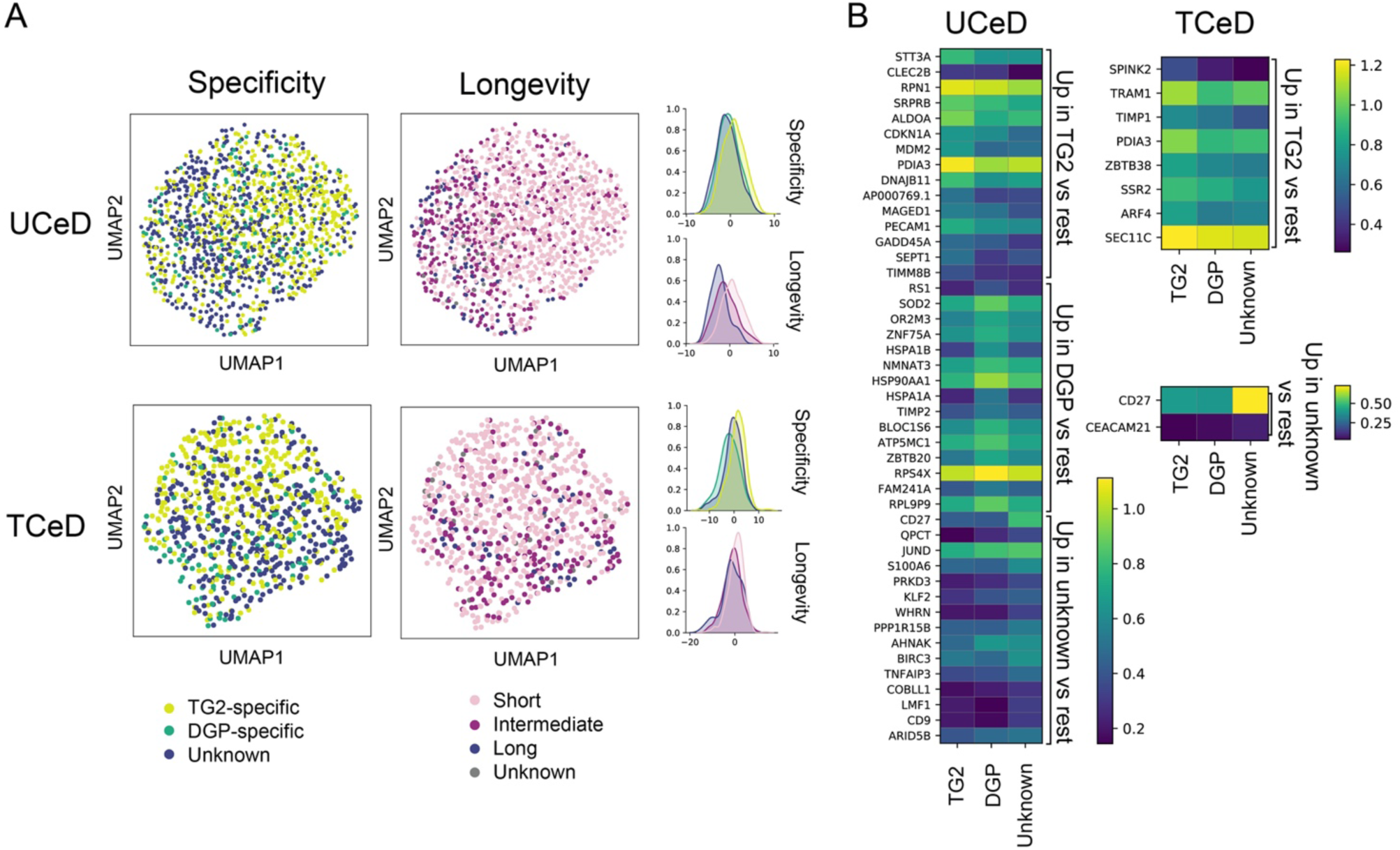
Transcriptional differences between TG2-specific PCs, DGP-specific PCs and PCs of unknown specificity in CeD. PCs from CeD patients were pooled together according to disease status (n=4 (UCeD), n=3 (TCeD)). Variation explained by patient (and sequencing library for TCeD) was regressed out, retaining variation explained by PC specificity. **A.** UMAP projections (left) of PCs from UCeD patients (top) or TCeD patients (bottom). UMAP is based on the first ten (UCeD) or five (TCeD) principal components. The individual principal components (PC1 for UCeD, PC2 for TCeD) with the most variation explained by specificity or longevity are shown (right). **B.** Heat maps showing the 15 most highly ranked statistically significant upregulated genes for each specificity group after filtering (*min_fold_change=1.2*).

In order to identify particular genes characteristic of the different specificities, we performed differential expression analysis for UCeD and TCeD patients separately (Fig. 5B). Strikingly, *CD27* was upregulated in PCs of unknown specificity compared to TG2- or DGP-specific PCs both in UCeD and TCeD. Genes involved in metabolism (*MDM2, PDIA3, SEPT1, STT3A, ALDOA*), regulation of cell cycle (*CDKN1A*), and the adhesion molecule CD31 (*PECAM1*) were upregulated among TG2-specific PCs in UCeD patients.

To investigate whether the transcriptional differences between PCs with different specificities could be explained by the skewed composition of longevity phenotypes, we repeated the differential expression analysis for cells of short longevity only. Notably, we found the majority of the upregulated genes in TG2-specific cells to be differentially expressed also when looking at short-lived PCs only (data not shown), indicating that the differential gene expression between TG2-specific and other PCs cannot be explained by differences in longevity phenotype only.

### PCs of CeD patients show altered expression of several genes

Having compared TG2- and DGP-specific PCs to PCs of unknown specificity in CeD, we next explored the possibility that intestinal PCs of CeD patients in general differ from PCs from healthy individuals. To test this hypothesis, we compared PCs of unknown specificity from CeD patients to PCs of controls. PCA revealed that principal component 3 (PC3) was the main driver of variation between controls, TCeD and UCeD patients, as visualized by UMAP (Fig. 6A). Longevity also explained some of the variation, but seemingly to a lesser degree than disease status.

**FIGURE 6.**
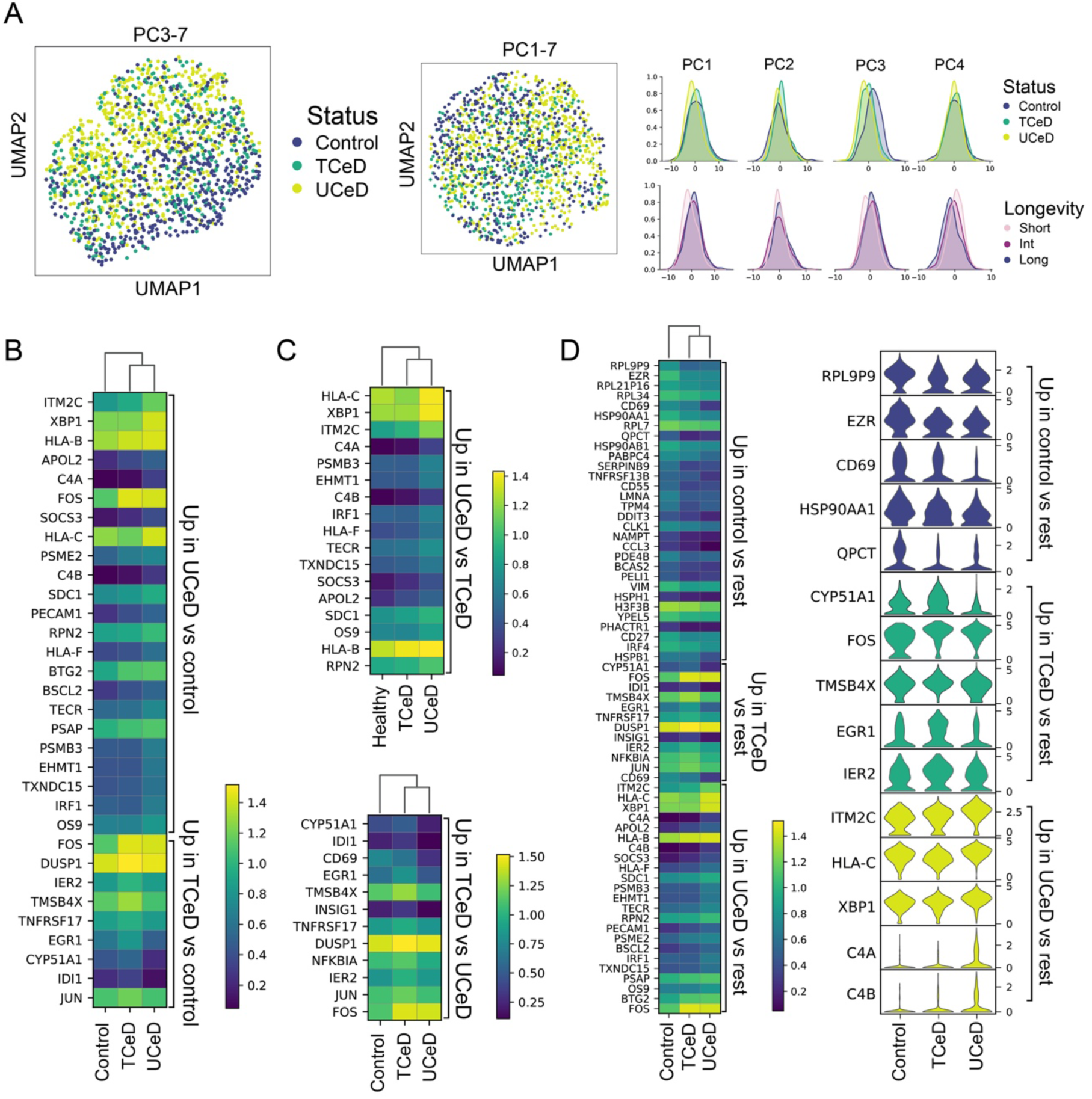
Transcriptional differences between PCs of unknown specificity in CeD patients and controls. PCs of unknown specificity from all UCeD patients (n=4), TCeD patients (n=3) and controls for which we sorted a natural composition of longevities (n=3) were pooled together. **A.** UMAP and individual principal component distributions of all PCs of unknown specificities. **B.** Heatmap of all differentially expressed (DE) genes between UCeD/TCeD and controls after filtering (*min_fold_change=1.3*). **C.** Heatmap of all DE genes between UCeD and TCeD (*min_fold_change=1.3*). **D.** Heatmap showing top 30 DE genes (*min_fold_change=1.3*) and violin plots showing top five DE genes (*min_fold_change=1.4*) between each group and the rest.

We then looked into individual genes that were differentially expressed between UCeD patients, TCeD patients and controls (Fig. 6B-D). Genes encoding complement proteins C4A and C4B, as well as *PECAM1* and *BSLC2*, were more highly expressed in CeD patients (in particular in UCeD) compared to the controls (Fig. 6B). Interestingly, we also observed some differentially expressed genes between UCeD and TCeD patients, including *XBP1, ITM2C, CD69, C4A* and *C4B*, with TCeD patients showing expression levels more similar to those of control subjects (Fig. 6C). Moreover, *CD69, QPCT, TNFRSF13B* (encoding TACI), *CCL3* and *CD27* were upregulated in controls compared to CeD patients (Fig. 6D). The majority of the genes that were differentially expressed between the disease statuses were also differentially expressed when looking only at short-lived cells (data not shown), demonstrating that the transcriptional differences between disease statuses are not mainly due to a skewed composition of cell longevities. Our single-cell analysis of intestinal PCs in CeD and controls thus indicates that intestinal PCs, in general, differ between CeD patients and healthy individuals.

### DGP-specific PCs derive from a few heavily expanded stereotypical clonotypes

In order to identify clonally related PCs of each specificity, we computationally reconstructed the paired heavy and light chain sequences of each BCR from our scRNA-seq data using BraCeR.^30^ We observed very little clonal expansion in the PC populations of unknown specificity, while there were many clone groups of moderate sizes for TG2-specific PCs and very large clonal expansions for the DGP-specific PCs (Fig. 7). Although the BCR repertoires of TG2-specific PCs in CeD have been extensively studied,^19,20,22^ fewer and more small-scale studies have been done for DGP-specific PCs.^10,21^

**FIGURE 7.**
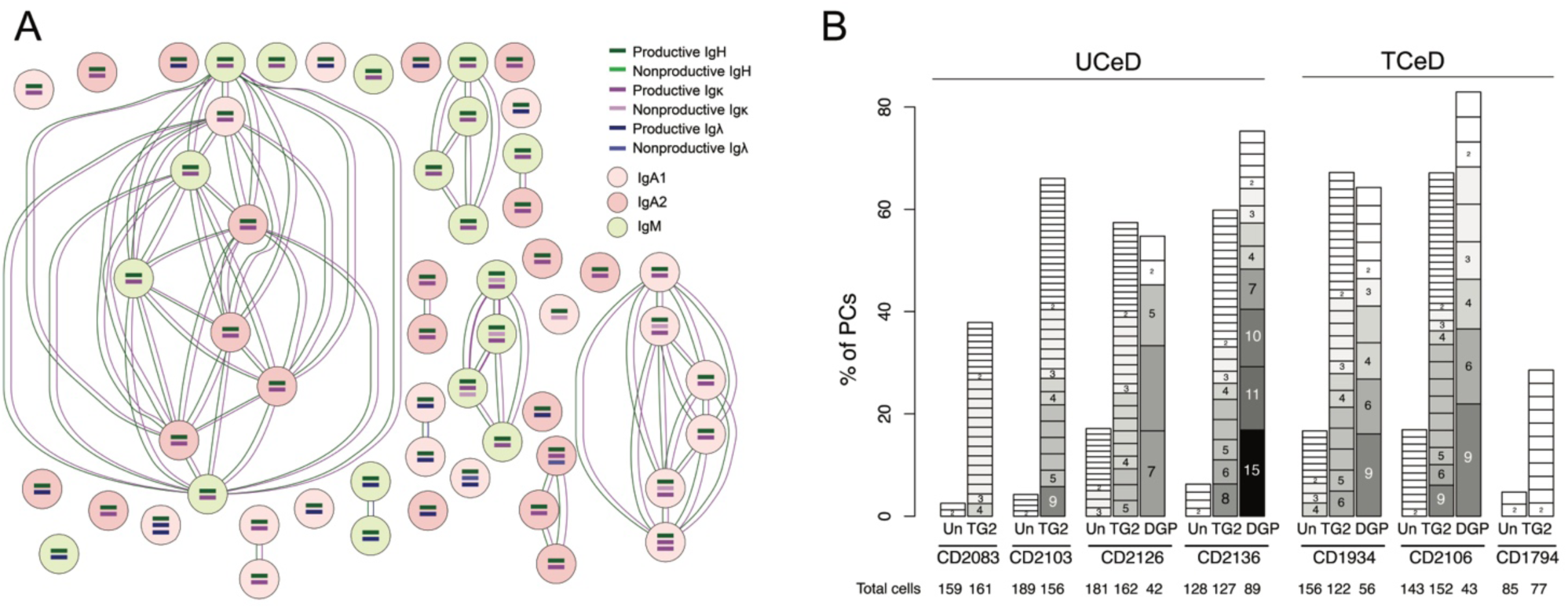
Clonal expansion of PCs in CeD. **A.** Example of clonal expansion of DGP-specific PCs from a TCeD patient (CD1934) as inferred by BraCeR. Circles represent individual cells, and connections indicate clonal relationship. **B.** Clone group sizes for each specificity in each CeD patient shown as percent of PCs. Darker shades of grey and larger boxes indicate bigger clones, with number of cells in each clone group shown. Only clones with two or more cells are shown. Un: Unknown specificity.

Here, by looking at the reconstructed BCR sequences of TG2-specific and DGP-specific PCs, we observed a similar skewing of variable (V)-gene usage as previously reported.^10,19-22^ TG2-specific PCs preferentially used *IGHV5-51, IGHV1-69* and *IGHV3-43*, with 66±3.9% of unique clones pairing with Igκ (in particular V-genes *IGKV1-5* and *IGKV3-11*). For DGP-specific PCs, we identified stereotypical V-gene pairing between *IGHV3-74* and *IGKV4-1*, which has not been previously reported.

### Importance of heavy chain residue R55, occasionally polymorphic, for binding to DGP

The novel stereotypical pairing of *IGHV3-74* and *IGKV4-1*, in addition to the previously reported *IGHV3-15*:*IGKV4-1* and *IGHV3-23(D)*:*IGLV4-69*,^21^ made up almost the entire repertoire of the DGP-specific PCs in all four CeD patients for which DGP-specific BCR sequences were available (Fig. 8A). Interestingly, *IGHV3-15* and *IGHV3-74* share a conserved arginine residue at IMGT position 55 (R55) (Fig. 8B). R55 is only present as a non-polymorphic residue in five *IGHV* genes, but occurs as an allelic variant of seven additional *IGHV* genes.^48^ R55 has previously been demonstrated to be essential for binding of DGP-specific antibodies using *IGHV3-15*:*IGKV4-1*.^21^ We therefore hypothesized that *IGHV3-74:IGKV4-1* antibodies could bind DGP in a similar manner to that of *IGHV3-15:IGKV4-1* antibodies.

**FIGURE 8.**
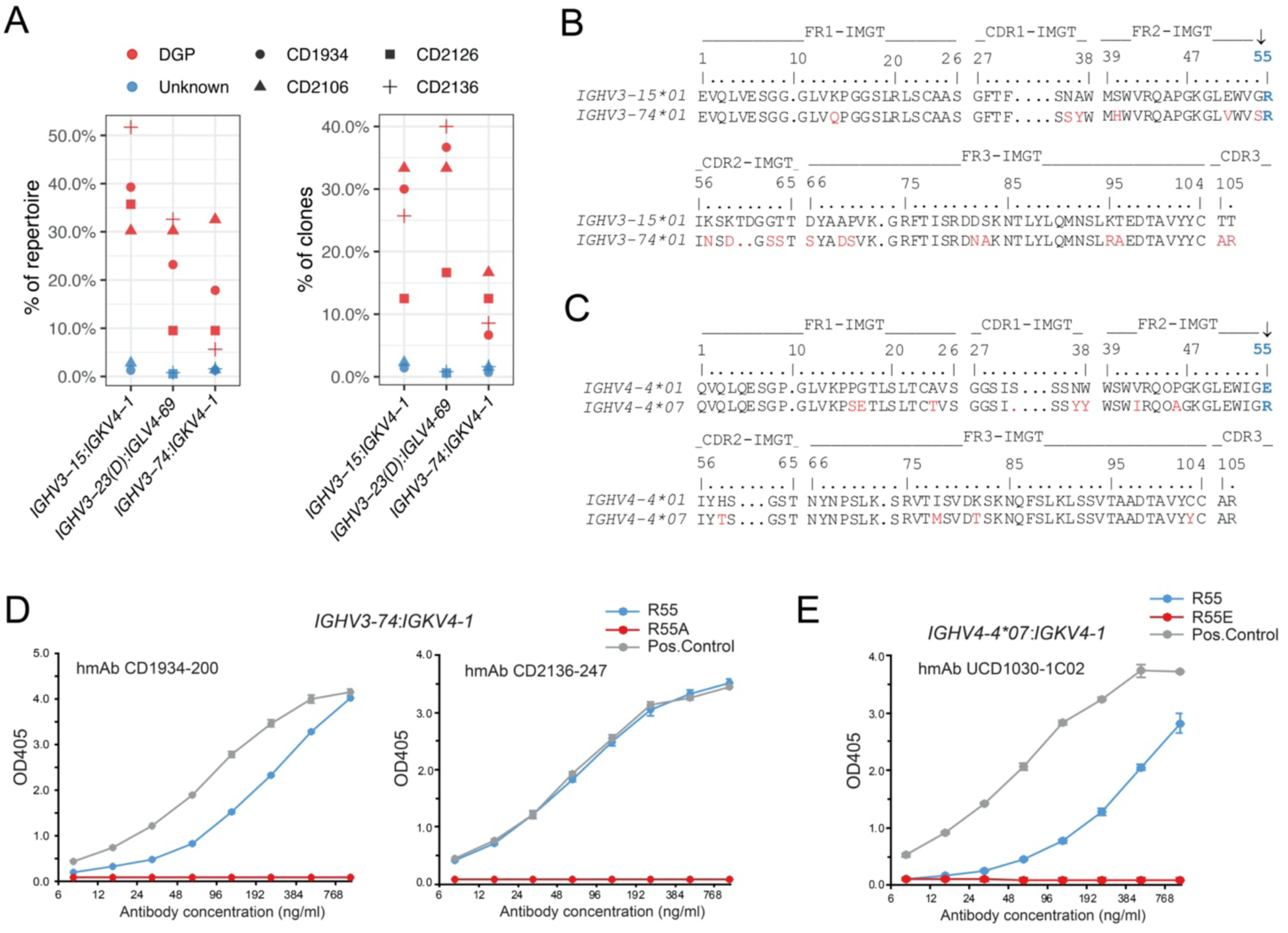
Repertoire analysis and site-directed mutagenesis of DGP-specific antibodies. **A.** Percent of total DGP-specific PCs (left) or DGP-specific clone groups (right) in each patient using the three dominating V-gene chain pairings. *IGHV3-23* and *IGHV3-23D* are collectively referred to as *IGHV3-23(D)*. **B.** Alignment of the amino acid sequences of *IGHV3-15* and *IGHV3-74* according to the IMGT numbering. The conserved R residue in position 55 is highlighted in blue and with an arrow. Positions that differ between the two gene segments are shown in red. **C.** Alignment of the amino acid sequences of *IGHV4-4*01* and *IGHV4-4*07* according to the IMGT numbering. The polymorphic R/E residue in position 55 is highlighted in blue and with an arrow. Other residue that differ between the two alleles are shown in red. **D, E.** Importance of heavy-chain R55 for DGP binding assessed by ELISA. Two patient-derived DGP-specific human monoclonal antibodies (hmAbs) using *IGHV3-74*:*IGKV4-1* reported in this study (D) and one previously reported DGP-specific hmAb (UCD1030-1C02)10 using *IGHV4-4*07*:*IGKV4-1* (E) were analyzed with or without R in position 55. A previously characterized DGP-specific hmAb (UCD1002-1E03)10 using *IGHV3-15*:*IGKV4-1* was used as positive control for comparison. Error bars indicate standard deviation based on sample duplicates.

To test the importance of R55 in the heavy chain of *IGHV3-74:IGKV4-1* antibodies for binding to DGP, we expressed two different human monoclonal antibodies (hmAbs) based on the BCR sequences of patient-derived DGP-specific PCs using the *IGHV3-74:IGKV4-1* chain pairing. We expressed each antibody in two variants with either R or A in position 55 of the heavy chain (R55 or R55A), and tested for reactivity to DGP in ELISA. While both hmAbs harboring R55 demonstrated reactivity to DGP, the R55A mutation led to a complete loss of DGP reactivity (Fig. 8D).

Interestingly, IMGT position 55 is polymorphic in *IGHV4-4*, with R55 present in the *\*07* allele and E55 in alleles \**01* through \**06* (Fig. 8C). Since immunoglobulin polymorphisms can influence antibody repertoires,^49^ we performed site-directed mutagenesis of R55 in a previously reported^10^ patient-derived DGP-specific hmAb using *IGHV4-4*07*:*IGKV4-1*, and showed that the R55E mutation disrupted binding to DGP (Fig. 8E).

### CeD-specific clonal lineage trees contain PCs spanning different isotypes and longevities

In addition to facilitating clonal inference and repertoire analysis, reconstruction of BCR sequences also allowed us to accurately determine the isotype of each PC. Inspection of individual clone groups and lineage trees revealed that several clone groups contained PCs of different isotypes and longevities (Fig. 9A-B).

**FIGURE 9.**
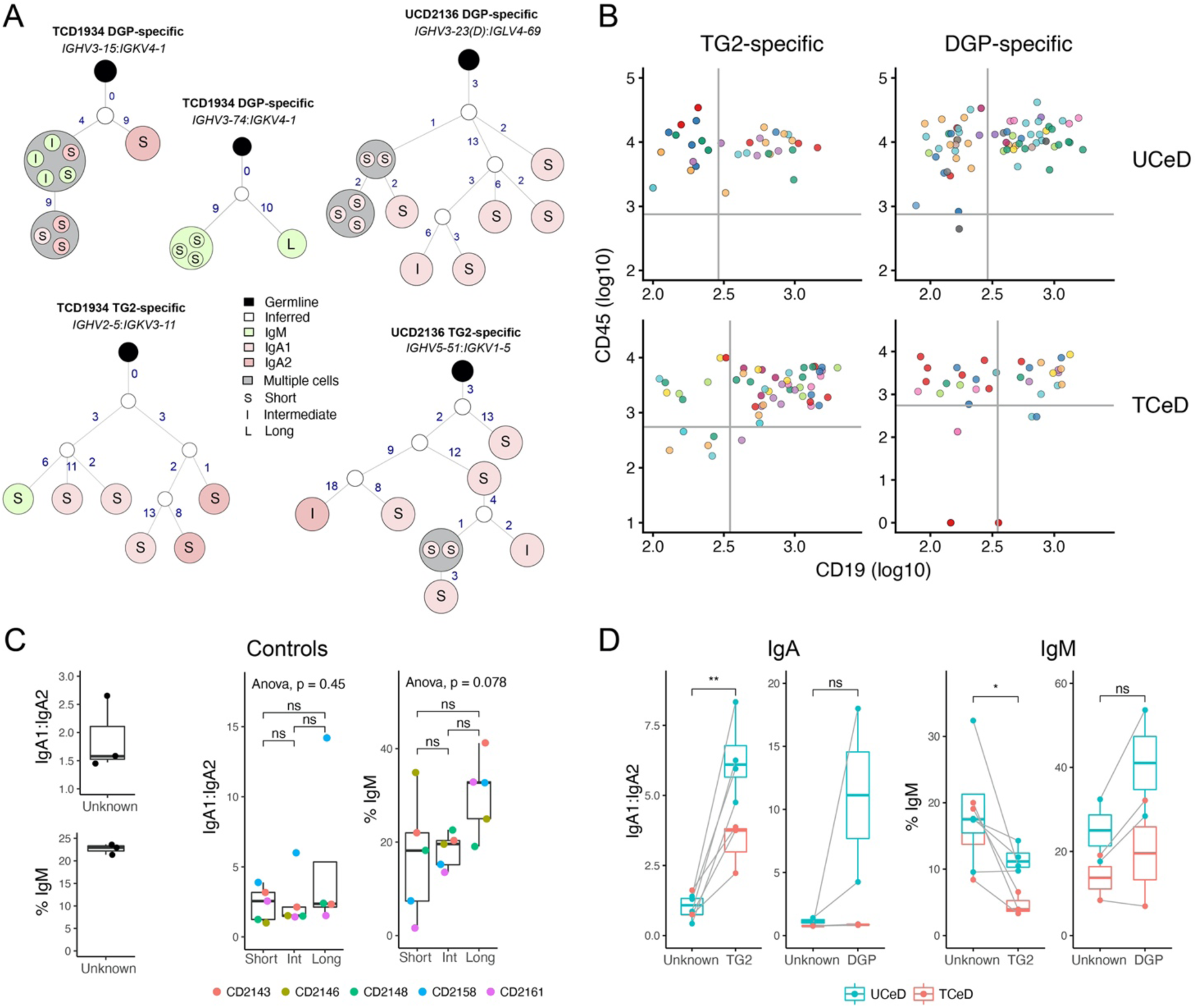
Clonality and isotype usage spanning different PC longevity populations. **A**. Example of lineage trees for DGP-or TG2-specific clones containing different isotypes and longevities from an untreated (UCD2136) and treated (TCD1934) patient. *IGHV3-23* and *IGHV3-23D* are grouped together as *IGHV3-23(D)*. **B.** Log10 index-sorting values for CD19 and CD45 for highlighted TG2-specific (left) and DGP-specific (right) clone groups for two representative patients (CD2136 and CD2106). Negative values were converted to 1 before they were logarithmized. Only the largest clones are shown, with members of each clone shown in the same color. **C.** IgA1:IgA2 ratio and IgM usage in controls (left). Only individuals for which a natural composition of PC longevities was sorted are shown. IgA1/IgA2 ratio (center) and percent IgM PCs (right) between PCs of different longevities in all controls are color coded according to patient. **D**. Skewed IgA subclass usage and IgM usage between disease-specific and other PCs in CeD.

It has long been known that IgA1 is preferentially used over IgA2 in the upper small intestine.^50^ When looking at isotype usage in the controls, we observed this preferred usage of IgA1 (Fig. 9C). The proportion of PCs expressing IgM was also in line with previous observations.^51^ While short-lived intestinal PCs have been shown to be skewed toward IgA usage,^5^ no reports of IgA subclass usage have been made. We therefore tested for differences in IgA1/IgA2 ratio between the three PC longevity populations in the controls, but found no statistically significant differences.

For both UCeD and TCeD, TG2-specific PCs had significantly lower usage of the IgM isotype (and higher usage of IgA) compared to PCs of unknown specificity from the same individuals (Fig. 9D). In line with previous observations, DGP-specific PCs seemed less restricted to the IgA-compartment than TG2-specific PCs^10^ (Fig. 9D). Notably, we also observed a significantly higher ratio of IgA1/IgA2 usage in TG2-specific PCs compared to PCs of unknown specificity, in line with a previous study.^22^

### SHM rates vary with isotype subclass, longevity, specificity and V-gene usage

Lastly, we looked into the SHM frequency in each reconstructed BCR sequence. In line with previous reports,^10,18-21^ TG2- and DGP-specific PCs had fewer somatic mutations compared to PCs of unknown specificity, and the number of mutations in heavy and light chains correlate (Fig. 10A). Notably, the DGP-specific PCs exhibited somewhat more mutations than the TG2-specific PCs. Unexpectedly, we also detected fewer somatic mutations in PCs of unknown specificity in UCeD patients compared to TCeD patients or controls (Fig. 10B).

**FIGURE 10.**
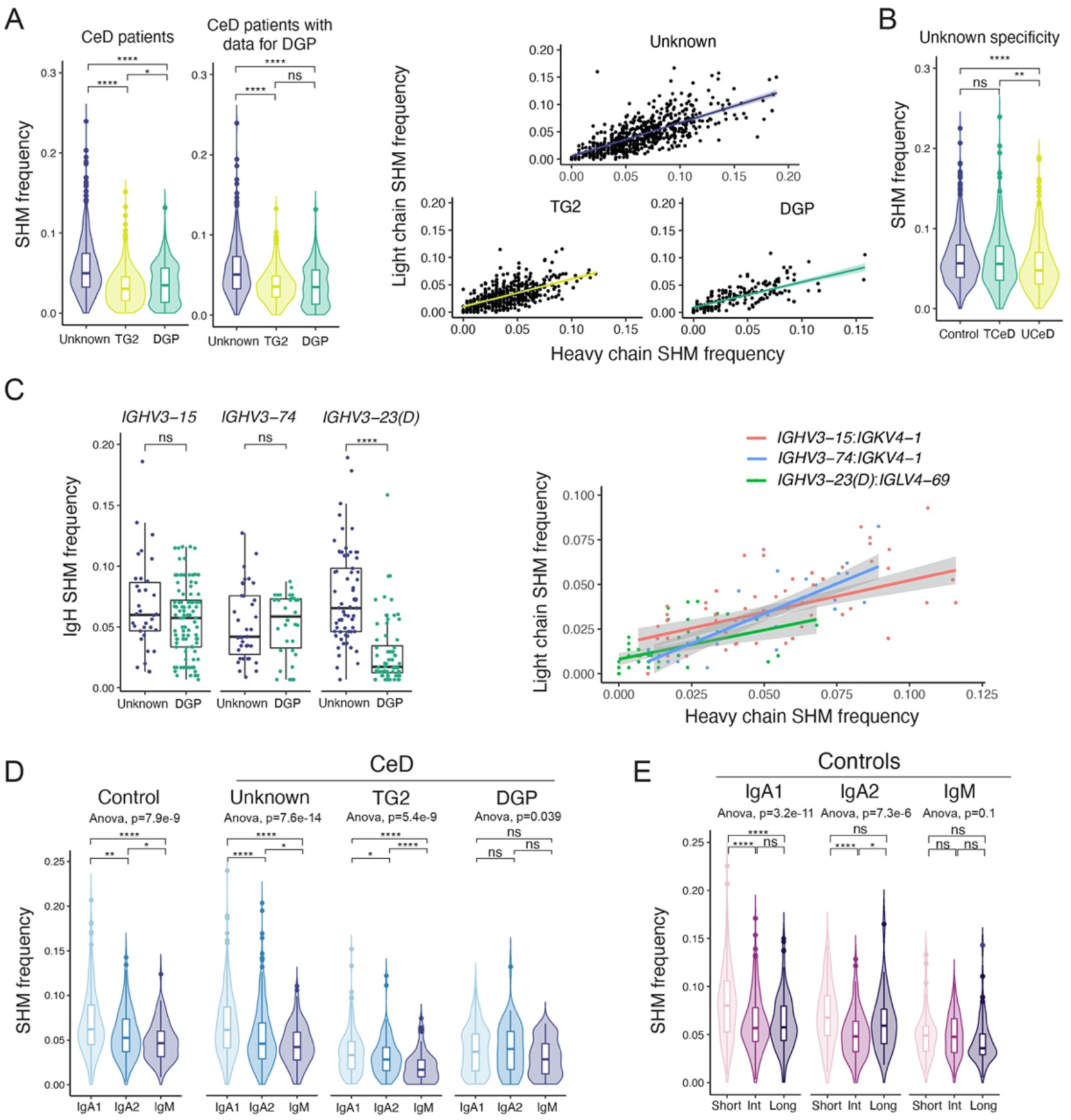
Somatic hypermutation frequency for PCs according to specificity, disease status, V-gene usage, isotype and longevity. Length-corrected rates of SHM in the V-region of reconstructed BCR sequences. Average SHM frequencies for heavy and light chain are shown, unless only one of the chains was reconstructed or otherwise stated. *p*-values were calculated with an unpaired Wilcoxon rank-sum test (*=*p*<.05, **=*p*<.01, ****=*p*<.0001). Adjusted *p*-values are shown. **A.** SHM frequency of all CeD patients (n=7) and patients for which DGP-specific PCs were sorted (n=4) stratified by specificity (left). Correlation between heavy and light chain SHM frequencies stratified by PC specificity are shown in the right panel for patients with sorted DGP-specific PCs (n=4). **B.** SHM frequency of PCs of unknown specificity for all patients and controls stratified by disease status. **C.** Heavy-chain (IgH) SHM frequency of DGP-specific PCs and PCs of unknown specificity from patients for which DGP-specific PCs were sorted (n=4), stratified by usage of the three stereotypical *IGHV* genes used by DGP-specific PCs (left). Correlation between heavy- and light-chain SHM frequencies in the three most common chain pairings among DGP-specific PCs are shown to the right. *IGHV3-23* and *IGHV3-23D* are collectively referred to as *IGHV3-23(D)*. **D.** SHM frequency of isotype (sub)classes from controls (left) and for each PC specificity in all CeD patients (right). Only cells from the three controls for which we did not artificially skew the composition of PC longevities are shown and were pooled together (n=3). **E.** SHM frequency of isotype subclasses and PC longevities from all controls (n=5).

DGP-specific PCs using *IGHV3-23(D)* have previously been shown to accumulate significantly fewer mutations than *IGHV3-23(D)* PCs of unknown specificity.^21^ We therefore investigated if this was the case also for DGP-specific PCs using *IGHV3-15*:*IGKV4-1* or *IGHV3-74*:*IGKV4-1*. In contrast to *IGHV3-23(D)*:*IGLV4-69* we found no significant differences in mutational load between DGP-specific PCs and PCs of unknown specificity using *IGHV3-15* or *IGHV3-74* (Fig. 10C, left panel). This finding indicates that the lower SHM frequency of DGP-specific PCs compared to PCs of unknown specificity is coupled to the usage of V-genes. Again, the frequencies of mutations in heavy and light chains were correlated (Fig. 10C, right panel).

We observed a higher rate of SHM in PCs of the IgA1 subclass compared to IgA2 and IgM in the controls, in line with what has previously been reported for peripheral blood mononuclear cells (PBMCs).^52^ This trend was also evident for PCs of unknown specificity and TG2-specific PCs in the CeD patients (Fig. 10D). We then investigated whether SHM rates differ between PC longevities. In order to remove bias in the form of different isotype preferences between the longevities, we looked at each isotype subclass separately. Interestingly, when looking at the most abundant isotype subclass, IgA1, short-lived PCs had significantly more mutations than intermediate- or long-lived PCs (Fig. 10E).

## DISCUSSION

This study provides information on the longevity profile of disease-specific PCs of CeD. We found that a substantial population of disease-specific PCs remained in patients even after at least one year on GFD, which is in keeping with a previous observation.^12^ Conceivably, this feature could imply that the cells which remain are mainly long-lived. This appears to be the case only in very long-term TCeD patients, as we demonstrate that in UCeD and in short-term TCeD patients most of the TG2- and DGP-specific PCs have the phenotype of short-lived CD19^+^CD45^+^ PCs, with a substantial population having an intermediate-lived CD19^-^CD45^+^ phenotype. This finding can explain why the TG2-specific PC population decreases already the first months after a patient starts on a GFD.^12^ While we found that the frequency of PCs with intermediate- and long-lived phenotype among the disease-specific PCs increased with time on GFD, the presence of disease-specific short-lived PCs even in long-term TCeD patients indicates that a significant proportion of such PCs must be continuously regenerated even on a GFD. A reason underlying this observation can be that most patients have occasional sporadic intake of small amounts of gluten.^53^ The disease-specific PCs with short-lived phenotype could potentially be formed by exposure to trace amounts of gluten. We also observe that PCs of different longevities exist within the same clonal lineages. This suggests that at least some of the short-lived PCs have been generated from memory B cells, as would fit with the observation of clonal relatedness between TG2-specific PCs and CD27^+^ memory B cells sampled from peripheral blood.^19^ Importantly, we observed a dominance of short-lived CD19^+^CD45^+^ phenotype in UCeD and short-term-treated CeD, both among disease-specific PCs but also in the general PC population. These data indicate that new and short-lived PCs massively accumulate in the lesion of UCeD and that the antigen targeting of many of these PCs is not directed to the disease-specific antigens.

Our transcriptional analysis of PC populations with different longevities revealed interesting differences. Short-lived PCs exhibited higher expression of a range of genes encoding ribosomal proteins and various genes involved in metabolism. Other genes involved in metabolism, *CD27*, and HLA class I genes were upregulated in long-lived PCs. These differences in gene expression may thus reflect previously reported differences in metabolic activity or antibody production between short- and long-lived PCs.^5,54^

Functional roles of PCs that go beyond antibody secretion have been suggested over the last years, including reports of cytokine production.^3^ Our results and our previously reported bulk RNA-seq data^15^ indicate that intestinal PCs produce mRNA transcripts for cytokines such as IL-15 and IL-16. IL-15 has been implicated in many autoimmune diseases and is a known player in CeD.^55^ It is not known, however, if all the cytokines for which we identified mRNA are actually translated and produced in the PCs. In the case of IL-15, for instance, it is known that its translation is under extensive regulation of a range of control elements, often resulting in repressed translation.^56-58^ Furthermore, two isoforms of IL-15 exist, one of which is not secreted but rather stored intracellularly.^57,59^

Conventionally, PCs are not thought to be APCs for T cells. However, the retained cell surface expression of a functional BCR in IgA- and IgM-expressing PCs^9,60,61^ make these cells able to fulfil the key hallmark of B cells as APCs having a cell surface receptor for antigen binding and uptake. In line with previously published bulk RNA-seq data^15^ we detected intermediate to low expression of HLA class II genes, in particular *HLA-DQA1*, and co-stimulatory molecules in a large fraction of the PCs. Indeed, a recent study found PCs to be the main cell type displaying DGP on disease-associated HLA class II molecules in the CeD lesion.^4^

We also showed, for the first time on a single-cell level, that TG2-specific PCs, DGP-specific PCs and PCs of unknown specificity display some transcriptional differences. A few of these differences can likely be ascribed to PC longevity. However, we also observed transcriptional differences between PC specificities when longevity was removed as a covariate suggesting that differences not related to potential longevity of the PCs may exist.

By comparing PCs of unknown specificity from CeD patients and controls, our work supports the recent finding that PCs of CeD patients generally may differ from PCs of healthy individuals.^15^ Notably, we also detected some differentially expressed genes between TCeD and UCeD patients, with TCeD patients often exhibiting intermediate expression values compared to UCeD or controls. Looking further into such differences may shed light on the potentially pathogenic roles of PCs in CeD, including the possibility that non-disease-specific PCs may also contribute to CeD.

PCs are generally thought to express high levels of CD27 on their surface. Interestingly, we found that *CD27* was upregulated both in long-lived versus short-lived PCs, PCs of unknown specificity versus TG2-specific and DGP-specific PCs, and in controls compared to CeD patients. Upregulation of *CD27* in non-TG2-specific compared to TG2-specific PCs has also been observed in bulk RNA-seq.^15^ Heterogeneous expression of *CD27* has previously been reported in multiple myeloma,^62^ and plasma blasts/PCs in patients with hemorrhagic fever with renal syndrome were found to have very low surface expression of CD27.^63^

We observed a somewhat higher frequency of TG2-specific PCs in UCeD and TCeD patients than previously reported.^10,18,19^ This discrepancy could be due to the fact that previous studies have focused on CD27^+^ IgA PCs, while we here made no such restrictions, and many of the TCeD patients in this study had been on a GFD for little more than one year. While we found only approximately 12% of TG2-specific PCs to be of the IgM isotype in UCeD, DGP-specific PCs seemed less restricted in their isotype usage, with only IgM PCs being detected in several clones. Unlike IgA antibodies, these IgM autoantibodies could be pathogenic through activation of complement via the classical pathway, as previously suggested.^64^

The preferential usage of IgA1 over IgA2 that we observed in PCs of the controls was already well-established.^50^ Beyond this known preference, we observed a further skewed IgA1/IgA2 usage in TG2-specific PCs compared to PCs of unknown specificity in line with a previous report from our group.^22^

While many clone groups contained only IgA1- or IgA2-expressing PCs, we also observed several clones spanning IgA1, IgA2 and even IgM. Although isotype class switching has been studied over the years,^65^ many details are still unclear. It has previously been suggested that switching to IgA2 could be a feature of the inducing antigen, as locations rich in bacteria often have a higher ratio of IgA2 cells.^66^ This seems unlikely to be the case for TG2 or DGP, as members of each clone group have been induced by the same antigen, but we still see isotype switch to both IgA subclasses in several clones.

IgA1 PBMCs are known to generally acquire more mutations than IgA2 or IgM PBMCs.^52^ Here we show that this is the case also for intestinal PCs. Moreover, we found that PCs of different longevities may acquire different numbers of mutations. When looking at IgA1 PCs, short-lived PCs had significantly more mutations compared to intermediate- or long-lived PCs in the controls. This is surprising, as TG2-specific PCs in CeD have strikingly low numbers of mutations^18-20^ despite being prone to more SHM due to their mainly short-lived IgA1 phenotype. This finding further emphasizes the functional importance of limited SHM in CeD-specific antibodies.

The finding of biased usage of *IGHV3-74*:*IGKV4-1* for recognition of DGP also underscores the importance of germline-encoded residues in CeD specific antibodies. Like in a previously identified stereotypic CeD antibody *IGHV3-15*:*IGKV4-1*,^21^ the *IGHV3-74*:*IGKV4-1* antibody utilizes heavy-chain residue R55 for recognition of DGP. This canonical function of R55, as we here demonstrate, was also observed for an *IGHV4-4*07*:*IGKV4-1* CeD antibody, where R55 is a polymorphic residue. While the latter is one antibody from one patient, and the overall impact of immunoglobulin polymorphisms for the formation of the CeD antibodies remains unexplored, this finding illustrates the principle that germline polymorphisms can indeed influence disease-specific antibody responses in CeD. More studies on the importance of immunoglobulin polymorphism for the humoral response in CeD are warranted.

## AUTHOR CONTRIBUTIONS

IL, CZ, LME, JP and S-WQ performed experiments. IL, CZ and ZM analyzed the data. KEAL and JJ provided patient samples. IL drafted the manuscript. LMS, RI, CZ, KEAL and S-WQ revised the manuscript. All authors approved the revised manuscript. LMS supervised the project.

## ACKNOWLEDGEMENTS

We would like to thank all patients donating samples, as well as C. Hinrichs, S. Furholm, M. Valde, S. Isaksen and F. van Megen for assistance with obtaining patient samples and clinical information. Further, we thank M.K. Johannesen, B. Simonsen and L. Wyrozemski for technical assistance with experiments. We also thank the Flow Cytometry Core Facility at OUS, in particular H. Notø, for assistance with flow cytometry, and R. Sandberg and G. Winberg at the Karolinska Institute for the generous gift of the plasmid encoding Tn5 transposase. Sequencing was performed at the Norwegian Sequencing Centre at Ullevål, Oslo, Norway, and raw sequencing data was stored and analyzed on the TSD (Tjeneste for Sensitive Data) facilities, owned by the University of Oslo. This work was funded by the University of Oslo World-leading research program on human immunology (WL-IMMUNOLOGY) and Stiftelsen KG Jebsen (project SKGJ-MED-017) to LMS, and Single Cell Gene Expression Atlas grant from the Wellcome Trust (108437/Z/15/Z) to ZM. Elements of Figure 1A were modified from Servier Medical Art, licensed under a Creative Commons Attribution 3.0 Unported License (http://smart.servier.com). The authors have no conflicting financial interests.

