## Supplementary Information for "Longevity, clonal relationship and transcriptional program of celiac disease-specific plasma cells"

#### SUPPLEMENTARY FIGURES

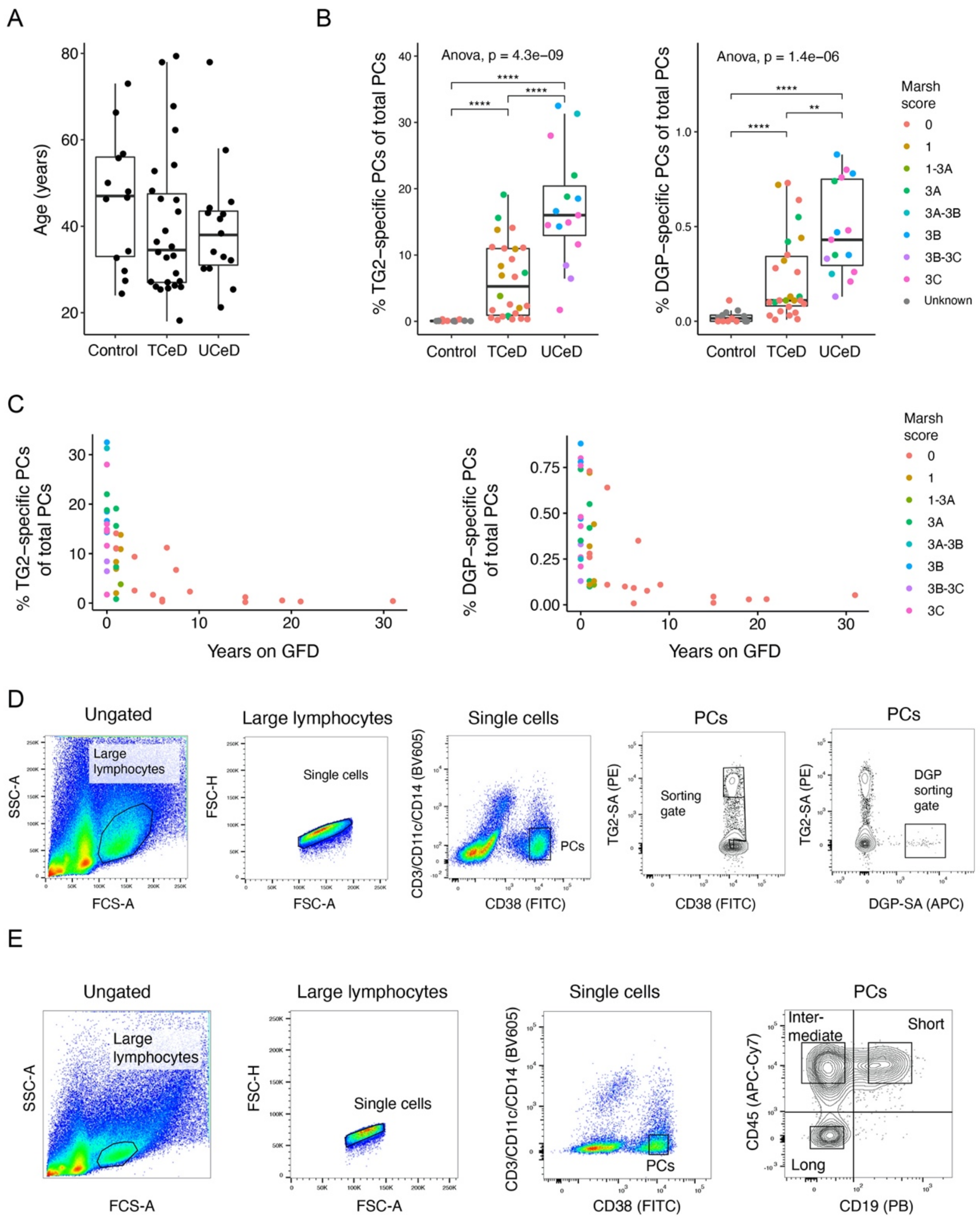


**SUPPLEMENTARY FIGURE 1. A.** Age distribution in patient groups. **B.** Percent of PCs identified as being specific for TG2 (left) or DGP (right) in UCeD (n=15), TCeD (n=26) and controls (n=13) by flow cytometry, colored according to the Marsh score of intestinal inflammation.[67] **C.** Percent of PCs identified as being specific for TG2 (left) or DGP (right) in UCeD (n=15) and TCeD patients (n=26) as a function of time on a gluten-free diet (GFD). **D.** Single-cell FACS index-sorting based on antigen specificity. Representative flow cytometry plots from one CeD patient are shown. **E.** Sorting based on CD45/CD19 surface marker expression as a proxy for cell longevity. Flow cytometry plots from one control are shown.

### SUPPLEMENTARY TABLES

**Supplementary Table 1:** Information on patients included in the study.

| *Subject* | *Status* | *Age* | *Gender* | *Marsh score at diagnosis*[a] | *Marsh score treated*[a] | *Years on GFD* | *Serum TG2-IgA*[b] | *Serum DGP-IgG*[c] | *HLA-DQ allotype* | *scRNA-seq data* |
|---|---|---|---|---|---|---|---|---|---|---|
| CD1541[d,e] | UCeD | 43 | M | 3B-3C | - | - | 27[f] | 4[f] | DQ2.5 | No |
| CD1742[e] | UCeD | 38 | F | 3B-3C | - | - | 14[f] | 5[f] | DQ2.5 | No |
| CD1993[g] | UCeD | 78 | M | 3C | - | - | 62.7 | 71 | DQ2.5 | No |
| CD2001 | UCeD | 46 | M | 3B | - | - | 33.7 | 18 | DQ2.5 | No |
| CD2002[g] | UCeD | 42 | F | 3B | - | - | 24.5 | >100 | DQ2.5 | No |
| CD2042[d,g] | UCeD | 25 | M | 3A | - | - | >100 | 80 | DQ2.5 | No |
| CD2079[g] | UCeD | 32 | F | 3B | - | - | 24.5 | >100 | DQ2.5 | No |
| CD2083 | UCeD | 30 | M | 3C | - | - | >100 | >100 | DQ2.5 | Yes |
| CD2103[g] | UCeD | 33 | F | 3C | - | - | 11.7 | >100 | DQ8 | Yes |
| CD2126 | UCeD | 30 | F | 3B | - | - | 70.5 | 99 | DQ2.5 | Yes |
| CD2136 | UCeD | 43 | F | 3C | - | - | >100 | 94 | DQ2.5 | Yes |
| CD2172 | UCeD | 44 | F | 3C | - | - | >100 | 56 | DQ8 | No |
| CD5012[e] | UCeD | 58 | M | 3A | - | - | 41[f] | 9[f] | DQ2.5 | No |
| CD5042[e] | UCeD | 34 | F | 3C | - | - | >128[f] | 61[f] | DQ2.5 | No |
| CD5044[e] | UCeD | 21 | F | 3A-3B | - | - | 76[f] | 30[f] | DQ2.5 | No |
| CD635[d] | TCeD | 68 | F | 3C | 0 | 15 | 2.8 | 5 | DQ2.5 | No |
| CD1305 | TCeD | 27 | F | 3B | 0 | 6.5 | <1.0 | 6 | DQ8 | No |
| CD1374[d] | TCeD | 27 | F | 3C | 0 | 3 | 1.2 | <5 | ND | No |
| CD1607 | TCeD | 33 | F | 3A | 1 | 1.5 | 3.6 | 15 | DQ2.5 | No |
| CD1618 | TCeD | 62 | F | 3A, 3C | 1-3A | 1.5 | 1.8 | 5 | DQ2.5 | No |
| CD1628[d] | TCeD | 29 | F | 3C | 1 | 1 | 2.1 | 5 | DQ2.5 | No |
| CD1650 | TCeD | 18 | F | 3B | 1 | 1.5 | 7.3 | 6 | DQ2.5 | No |
| CD1794 | TCeD | 26 | M | 3C | 0 | 1 | <1.0 | 8 | DQ2.5 | Yes |
| CD1934 | TCeD | 26 | F | 3C | 3A | 1 | 9.7 | 29 | DQ2.5 | Yes |
| CD1945 | TCeD | 78 | F | 3B | 1 | 1 | 2.0 | <5 | DQ2.5 | No |
| CD1992 | TCeD | 35 | F | 3C | 3A | 1 | 4.8 | 11 | DQ2.5 | No |
| CD1993[g] | TCeD | 79 | M | 3C | 3A | 1 | 5.4 | 12 | DQ2.5 | No |
| CD2002[g] | TCeD | 43 | F | 3B | 0 | 1 | 2.2 | 20 | DQ2.5 | No |
| CD2014[d] | TCeD | 54 | F | 3A | 0 | 31 | <1.0 | <5 | DQ2.5 | No |
| CD2042[d,g] | TCeD | 26 | M | 3A | 3A | 1 | >100 | 55 | DQ2.5 | No |
| CD2047[d] | TCeD | 25 | F | 3A | 0 | 7.5 | 2.9 | <5 | DQ2.5 | No |
| CD2049[d] | TCeD | 48 | F | 3A, 3B | 0 | 5 | <1.0 | <5 | DQ2.5 | No |
| CD2050[d] | TCeD | 36 | F | 1[h] | 0 | 6 | <1.0 | <5 | DQ8 | No |
| CD2057[d] | TCeD | 26 | F | 3B | 0 | 15 | <1.0 | <5 | DQ2.5 | No |
| CD2067[d] | TCeD | 46 | F | 3B, 3C | 0 | 6 | <1.0 | <5 | DQ2.5 | No |
| CD2075[d] | TCeD | 39 | F | 3B | 0 | 21 | 1.7 | <5 | DQ2.5 | No |
| CD2079[d,g] | TCeD | 33 | F | 3B | 0 | 1 | 7.5 | 7 | DQ2.5 | No |
| CD2081[d] | TCeD | 27 | F | 3 | 0 | 9 | 1.1 | <5 | DQ8 | No |
| CD2097[d] | TCeD | 53 | F | 3B | 0 | 19 | 1.8 | 5 | DQ2.5 | No |
| CD2103[d,g] | TCeD | 34 | F | 3C | 1 | 1 | 2.7 | 8 | DQ8 | No |
| CD2106 | TCeD | 46 | F | 3B-3C | 0 | 3 | 2.3 | 8 | DQ2.5 | Yes |
| CD1740 | Control | 47 | F | 0 | - | - | <1.0 | <5 | DQ7.5 | No |
| CD1741 | Control | 56 | F | 0 | - | - | <1.0 | <5 | DQ8 | No |
| CD1928 | Control | 24 | F | 0 | - | - | <1.0 | <5 | DQ2.5 | No |
| CD1947 | Control | 66 | M | ND | - | - | ND | ND | ND | No |
| CD2143 | Control | 27 | F | 0 | - | - | <1.0 | <5 | DQ2.5 | Yes |
| CD2145 | Control | 57 | F | 0 | - | - | ND | ND | ND | No |
| CD2146 | Control | 30 | M | 0 | - | - | ND | ND | ND | Yes |
| CD2147 | Control | 48 | M | ND | - | - | ND | ND | ND | No |
| CD2148 | Control | 33 | M | ND | - | - | ND | ND | ND | Yes |
| CD2158 | Control | 34 | M | 0 | - | - | <1.0 | <5 | DQ2.5 | Yes |
| CD2161 | Control | 46 | M | ND | - | - | <1.0 | <5 | DQ8 | Yes |
| CD2162 | Control | 50 | F | ND | - | - | <1.0 | <5 | DQ2-/DQ8- | No |
| CD2164 | Control | 73 | F | ND | - | - | <1.0 | <5 | DQ7.5 | No |

[a] Biopsies were scored according to the modified Marsh criteria as published by Oberhuber *et al*.[67]

[b] Measured with assay from Inova Diagnostics, with upper limit of normal 4 unless otherwise indicated.

[c] Measured with assay from Inova Diagnostics, with upper limit of normal 20 unless otherwise indicated.

[d] Sample has been cryopreserved.

[e] Samples from Akershus University Hospital.

[f] Measured with assay from Thermo Fisher, with upper limit of normal value of 7.

[g] Same patient analyzed both as UCeD and TCeD.

[h] Probable CeD according to British Society of Gastroenterology guidelines.[34]

M: male, F: female, UCeD: untreated celiac disease, TCeD: treated celiac disease, GFD: gluten-free diet, TG2: transglutaminase 2, DGP: deamidated gliadin peptide, HLA: human leukocyte antigen, scRNA-seq: single-cell RNA sequencing, ND: not determined.

**Supplementary Table 2:** Longevity analysis by flow cytometry based on the cell surface markers CD19 and CD45. Longevity of DGP-specific PCs is only shown for patients with more than 30 recorded DGP-specific PCs. (Excel file)

**Supplementary Table 3:** Library-specific quality control threshold values and cell numbers for scRNA-seq data.

| **Library** | **Subject** | **Diagnosis** | **Reads ($10^5$)** | | **Genes** | | **# Cells** | | **Specificity (# passing QC)** | | | **Longevity (# passing QC)** | | |
|---|---|---|---|---|---|---|---|---|---|---|---|---|---|---|
| | | | **Min** | **Max** | **Min** | **Max** | **Total** | **Pass** | **TG2** | **DGP** | **Unknown** | **Short** | **Int** | **Long** |
| **scLib3** | CD2083 | UCeD | 2.8 | 13.0 | 1200 | 4400 | 376 | 320 | 161 | 0 | 159 | 169 | 116 | 30 |
| **scLib4** | CD2103 | UCeD | 3.0 | 13.0 | 2000 | 4700 | 373 | 346 | 156 | 1 | 189 | 231 | 79 | 27 |
| **scLib5** | CD2106 | TCeD | 2.2 | 11.5 | 2000 | 4900 | 374 | 338 | 152 | 43 | 143 | 190 | 88 | 41 |
| **scLib6** | CD2126 | UCeD | 2.0 | 17.0 | 1200 | 4900 | 371 | 325 | 162 | 2 | 161 | 252 | 54 | 10 |
| **scLib7** | CD2136 | UCeD | 2.0 | 15.5 | 2000 | 4400 | 373 | 344 | 127 | 89 | 128 | 201 | 130 | 8 |
| **scLib8** | CD1934 | TCeD | 2.0 | 14.0 | 1300 | 4400 | 376 | 334 | 122 | 56 | 156 | 274 | 50 | 8 |
| **scLib9** | CD2143 | Control | 2.0 | 16.0 | 1900 | 4300 | 188 | 174 | 1 | 1 | 172 | 93 | 60 | 17 |
| | CD2146 | | | | | | 187 | 157 | 1 | 0 | 156 | 43 | 101 | 9 |
| **scLib10** | CD2148 | Control | 2.0 | 16.0 | 1500 | 4000 | 188 | 175 | 0 | 0 | 175 | 44 | 108 | 21 |
| | CD1794 | TCeD | | | | | 185 | 162 | 77 | 0 | 85 | 97 | 65 | 0 |
| **scLib11** | CD2158 | Control | 2.0 | 15.0 | 1300 | 3500 | 186 | 169 | 0 | 0 | 169 | 56 | 58 | 55 |
| | CD2161 | | | | | | 188 | 167 | 0 | 0 | 167 | 56 | 54 | 57 |
| **scLib12** | CD2158 | Control | 2.0 | 18.0 | 1200 | 4400 | 188 | 139 | 0 | 0 | 139 | 39 | 41 | 59 |
| | CD2161 | | | | | | 94 | 41 | 0 | 0 | 41 | 11 | 7 | 23 |
| | CD2126 | UCeD | | | | | 96 | 60 | 0 | 40 | 20 | 46 | 12 | 2 |
| **Total** | | | | | | | **3743** | **3251** | **959** | **232** | **2060** | **1802** | **1023** | **367** |

UCeD: untreated celiac disease, TCeD: treated celiac disease, QC: quality control, TG2: transglutaminase 2, DGP: deamidated gliadin peptide, Short: short-lived phenotype ($CD19^+CD45^+$), Int: intermediate-lived phenotype ($CD19^-CD45^+$), Long: long-lived phenotype ($CD19^-CD45^-$).
